# APPLICATION OF PLANTS BASED MATERIAL FOR THE MANUFACTURING OF MOSQUITO REPELLENT SURFACE COATING

**DOI:** 10.1101/2023.04.03.535318

**Authors:** Tehreema Iftikhar, Hammad Majeed, Rida Abid, Faheem. A. Khan

## Abstract

Mosquitos have been a malice and source of many diseases in humans. Later on, humans understood they learned that plants also possess mosquito-repellent properties. Different insectrepellent coatings are present in the market which are chemically prepared and can be harmful to humans and the environment. Different plants have insect-repellent properties which have been utilized in this research to make a nature-based insect-repellent surface coating. *Moringa oleifera* L. and *Mentha piperita* L. are naturally insect-repellent plants. Nanoparticles increase the surface area and efficiency of extracts of plants. Thus, ZnONP of *Moringa oliefera* L. and *Mentha piperita* L. plants were made characterization was done through UV-vis spectroscopy, FTIR, and PSA. The UV-visible spectrum showed absorption peaks for ZnO nanoparticles at 350nm for *Mentha piperita* L. and 356nm for *Moringa oleifera* L. The particle size analysis indicated the variable sizes of ZnONPs for both plants. FTIR showed vibration peaks from 3341 to 650cm^-1^ for *Moringa oleifera* L. and 3393 to 700 cm^-1^ for *Mentha piperita* L. ZnONPs were used in paint along with water extracts of plants to make the paint insect-repellent in nature. Mosquito repellent activity of paint formulations was also tested against *Aedes aegypti*.

## Introduction

Vector-borne diseases are caused by infections spread to humans and animals by blood-consuming arthropods such as mosquitoes, ticks, fleas, etc. Vector-borne diseases such as Dengue, Malaria, Chikungunya, and Yellow fever, have spread to almost all continents in past decades due to certain climatic conditions including Asia [1]. Mosquitoes have accomplished to be carriers for many infectious diseases which influence millions of people and result in a huge number of deaths. Therefore, it is essential to develop methods to prevent them. The control measures in use focus on a small group of mosquitoes by using chemical-based solutions such as coils and mats. The use of synthetic mosquito repellents may cause allergic reactions in some people [2]. Synthetic repellents can be very expensive and can harm the environment. Thus, it is extremely important to work on nature-based mosquito repellent.

A number of mosquito repellent paints and surface coatings are present in markets which are produced by a number of paint-producing companies such as MozziGuard by Nippon paint, Anti-Mosq by Kansai paint, Mosquito repellent paint by Plascon and Corion repel by Corion paints [3]. But the drawback is that all of these paints are synthetic with harmful chemicals to kill or repel mosquitoes and other insects. These chemicals are non-biodegradable and cause environmental pollution and they also affect living beings other than insects. These can also be harmful to humans as they can cause allergies to them which can be fatal if not treated quickly [4].

Essential oils which belong to several species of plants possess combinations of hydrocarbons that are found as an effective repellent against many insects. It has been discovered that the monoterpenoids, which make up the majority of the component, are cytotoxic to animal and plant tissue, compromising the proper operation of affected tissues [5].The plants used in this research are *Moringa oleifera Lam*. and *Mentha piperita* L. as these plants possess natural insect-repellent components.

## Methodology

### Extraction of Plant extracts

Microwave-assisted extraction was carried out to obtain aqueous plant extracts. 10g of plant powder was taken in 100ml dis. water and heat for 5 mins in a microwave oven with an interval of 30 sec after each minute. It was filtered and stored for further use [6].

### Preparation and characterization of Nanoparticles with Plant extracts

1mM solution of zinc sulfate was prepared by mixing 1.4 g of ZnSO4 salt in 50 ml distilled water [7]. ZnO nanoparticles were prepared using water extracts of *Moringa oliefera* and *Mentha piperita*. 50 ml of ZnSO4 salt solution was placed on a magnetic stirrer at 1200 rpm and plant extracts were added dropwise to the salt solution. Then plant extracts were added in it dropwise, 40ml of each extract. Color changed from golden yellow to dark bluish and greenish brown for *Moringa oleifera* and *Mentha piperita* respectively, this color change indicated the formation of ZnO nanoparticles for each plant specimen. The solution was stirred for 60 minutes on a magnetic stirrer. Then the solution was transferred to 50ml falcon tubes and centrifuged at 4000 rpm for 30 mins. The supernatant was discarded and pallets were transferred into Petri plates and dried at room temperature which was later finely powdered and stored at room temperature [7].

### Characterization of zinc oxide (ZnO) Nanoparticles

Characterization of ZnO nanoparticles was done through UV-Visible Spectroscopy, Particle Size Analysis (PSA) and Fourier transform infrared (FT-IR) spectroscopy [8] [9].

### Paint formulations using Plant-based additives

Aqueous extracts and ZnO particles of plants were taken to Diamond Paints Industry which is located in Sundar Industrial Zone, Lahore. Paint formulations were made by adding our plant-based material into water-based Plastic emulsion paint. We used plain white emulsion paint which did not have any preservative, biocide, and color in it so we can assess what effects or changes occurred after adding the plant-based material. A 3% ratio was used for adding plant material in each 200 ml paint sample. 0.6g ZnO particles of each Moringa and Peppermint were added in 200ml emulsion paint samples in a plastic jar and thoroughly mixed and then sealed for further use. 6ml plant extract of each Moringa and Peppermint was added in 200ml emulsion paint jars. In one sample standard biocide was added in the same ratio to compare the results of plant-based additives and one sample was taken as a control.

**Table 1:** Sample coding.

| Sample | Code |
| --- | --- |
| Paint with Moringa NPs | Sample 5 |
| Paint with Moringa Extract | Sample 6 |
| Paint with Peppermint NPs | Sample 7 |
| Paint with Peppermint Extract | Sample 8 |
| Paint with biocide | Standard biocide |
| Paint without biocide | Without biocide |

### The whiteness index assessment of Paint Formulation

The sheets of paint with plant-based additives were inspected for their whiteness. The sheets were observed under a reflectance Spectrophotometer. The change in color was noted after the addition of plant-based additives. The change in color was compared with CIE standard illuminant D65 at a temperature of 50 °C [10].

#### Mosquito repellent activity of Paint formulation

A mosquito repellency test was performed following [2] with some alternations. *Aedes aegypti* adult mosquitoes were provided by the insectary of the Department of Zoology GCU, Lahore. About 3-4 days old mosquitoes were reared at room temperature (28°C). The analysis was conducted in a transparent glass box measuring 60cm x 44cm x 29cm at ambient conditions (30°C, 50% humidity). Six test panels with the size of 14cm x 8cm, four were painted with paint with plant-based additives and one with biocide paint, and one without biocide which was considered as control. About 10 mosquitoes were released into the box for each test. The number of mosquitos landing on test panels was recorded every 5-minute intervals for up to 2 hour time. The rate, number, and frequency of the mosquitoes approaching the panels were recorded manually.

## Results and Discussion

### Synthesis and characterization of ZnO Nanoparticles of Plant extracts

In a 50ml solution of ZnSO4, 40ml plant extract was added slowly with constant stirring at 1200rpm on a Magnetic stirrer. The color change was observed which indicated the formation of nanoparticles. The color changed from dark yellow to blue-greenish for Peppermint and dark brown for Moringa. Characterization of ZnO nanoparticles was done through UV-Visible Spectroscopy, Particle Size Analysis (PSA) and Fourier transform infrared (FR-IR) spectroscopy [7].

UV-Visible spectroscopy was utilized to check the creation of Zinc Oxide nanoparticles. The UV-visible spectrum showed absorption peaks for ZnO nanoparticles at 350nm for *Mentha piperita* L. and 356nm for *Moringa oleifera* L. (Graph 1 a&b), which confirmed the formation of ZnO. The size of ZnO nanoparticles was determined using Particle Size Analyzer (Cary 360). 1mg of powder was mixed in 10ml of distilled water using a vortex mixer. PSA showed particles ranging from nano to fine in size. The size of Moringa ZnO particles was 27-614nm and that of Peppermint ZnO particles was 367-838nm (Graph 2 a&b). Fourier transform infrared (FR-IR) spectroscopy showed which indicated the occurrence of functional groups assisting the formation of nanoparticles (Graph 3 a&b). FT-IR showed the vibration peaks for ZnONPs are in the range of 600 cm^-1^ to 800 cm^-1^.

**Table 2:**
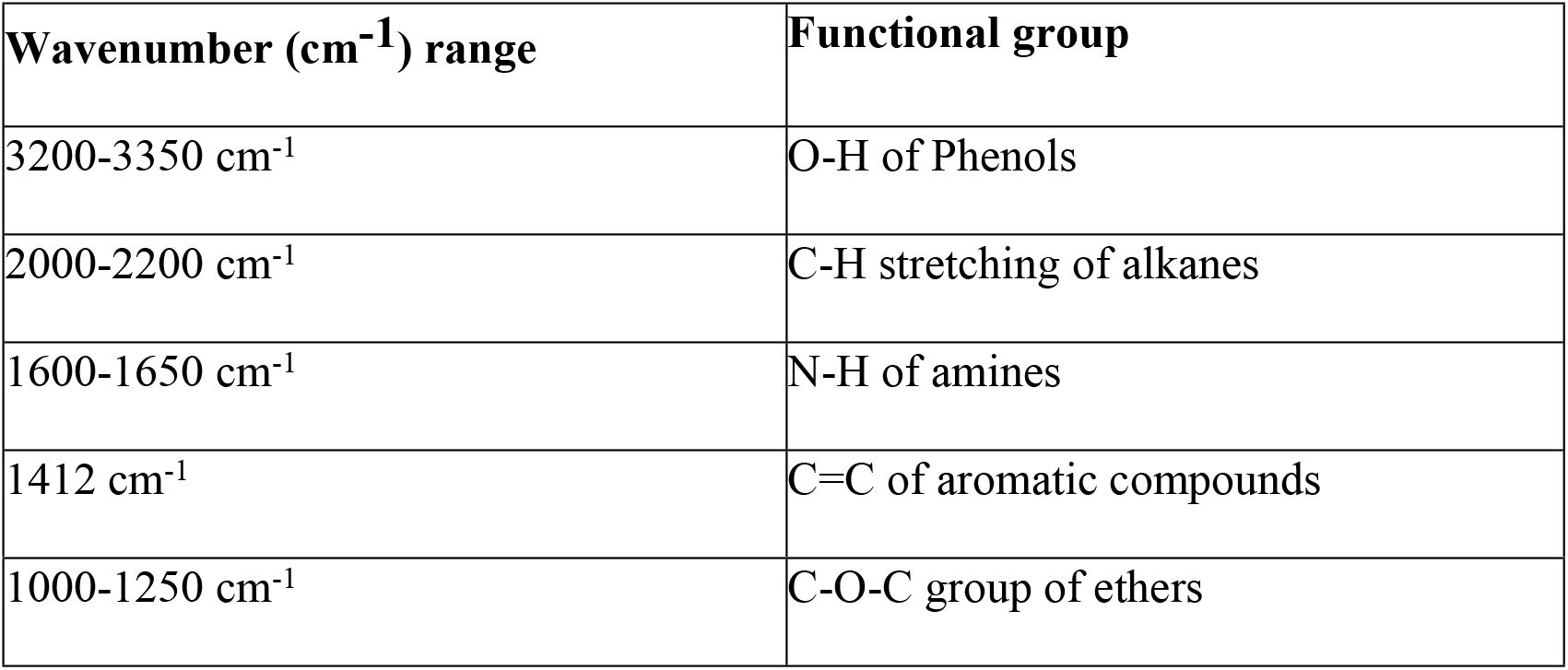
Different functional group peaks in the FTIR spectrum.

**Graph 1 (a):**
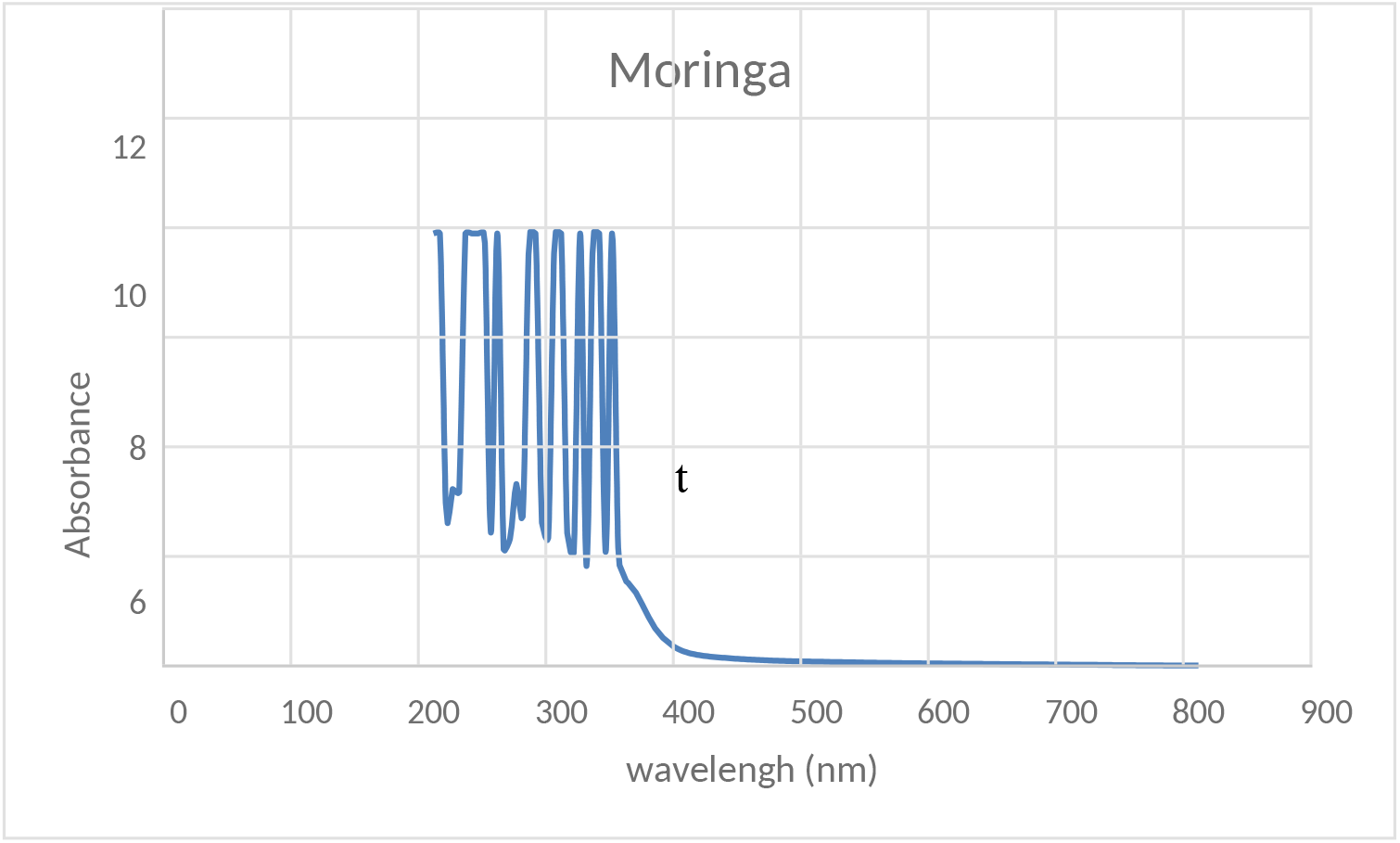
UV-Visible spectroscopy of *Moringa oleifera* ZnO nanoparticles.

**Graph 1 (b):**
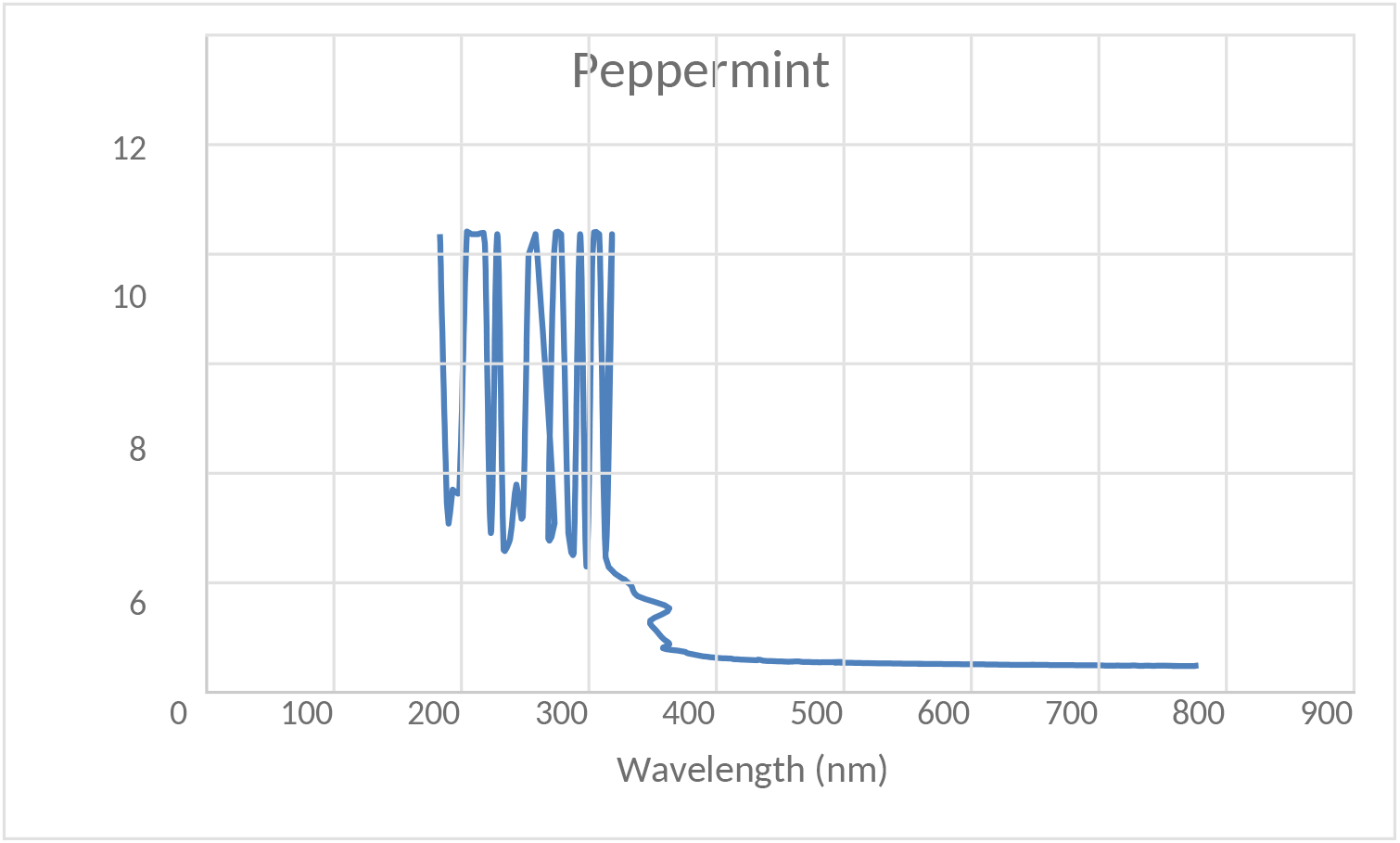
UV-Visible spectroscopy of *Mentha piperita* ZnO nanoparticles.

**Graph 2(a):**
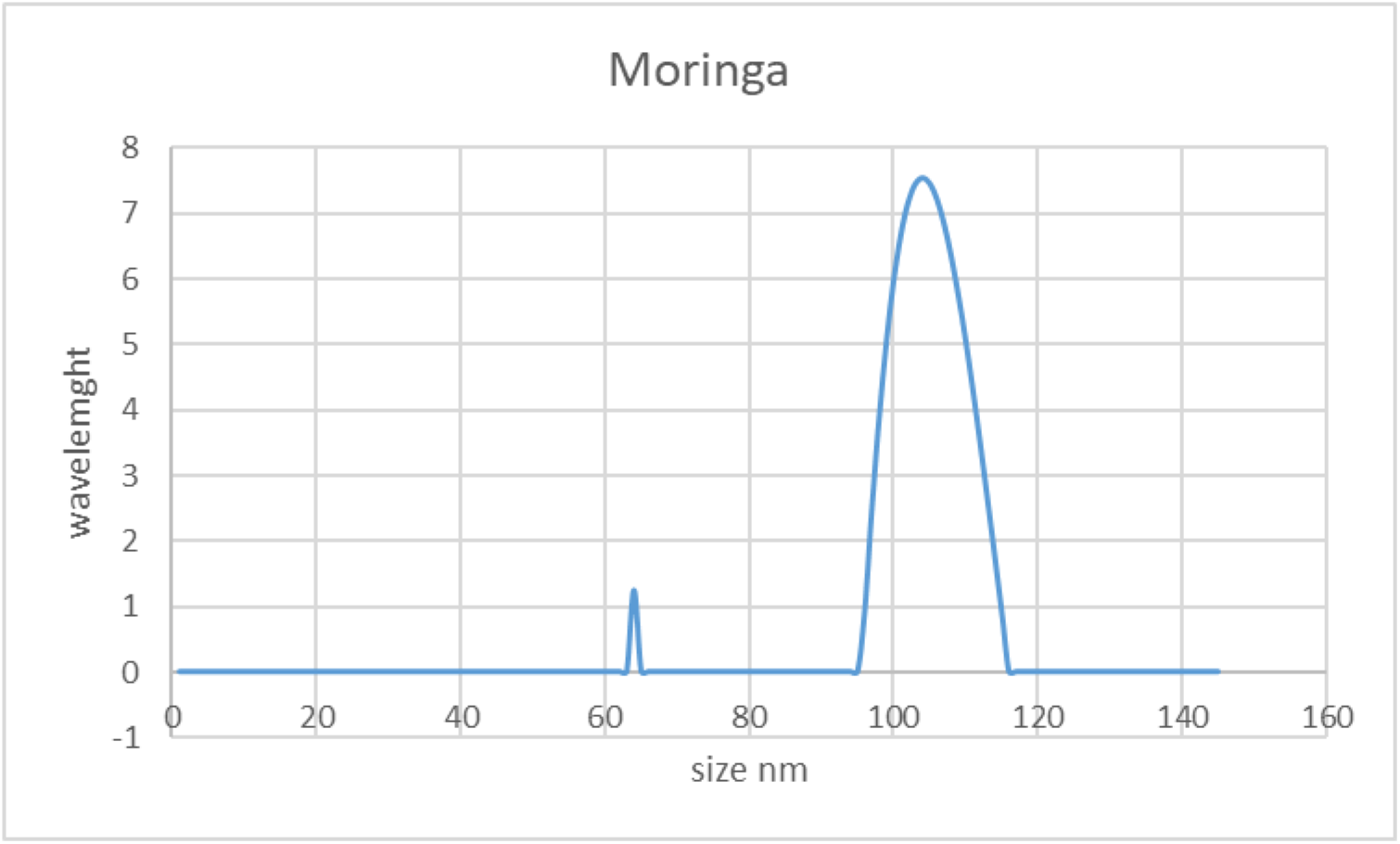
Particle size analysis of *Moringa oleifera* ZnO nanoparticles.

**Graph 2(b):**
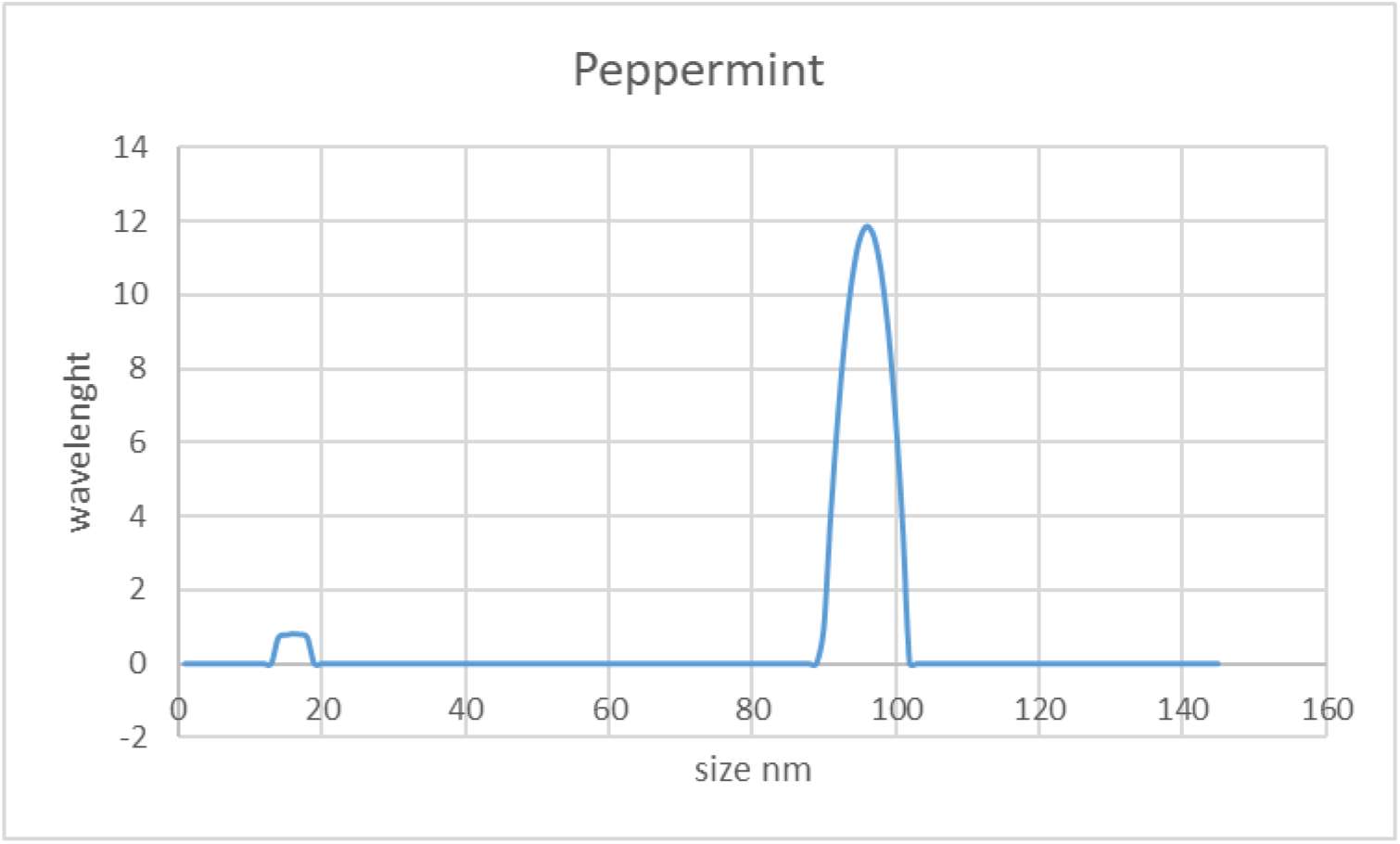
Particle size analysis of *Mentha piperita* ZnO nanoparticles.

**Graph 3 (a).**
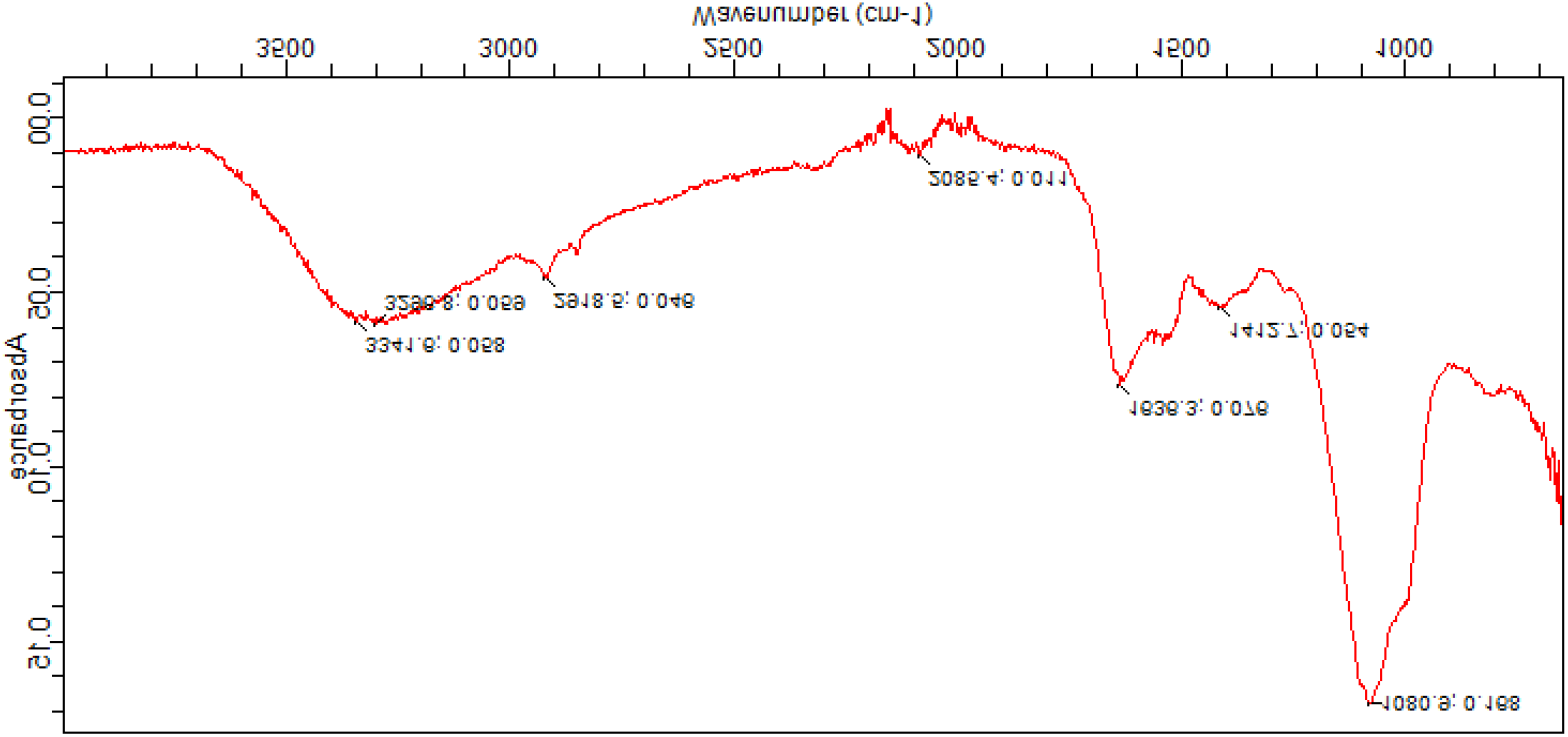
FT-IR spectrum of *Moringa oleifera* ZnO nanoparticles.

**Graph 3 (b).**
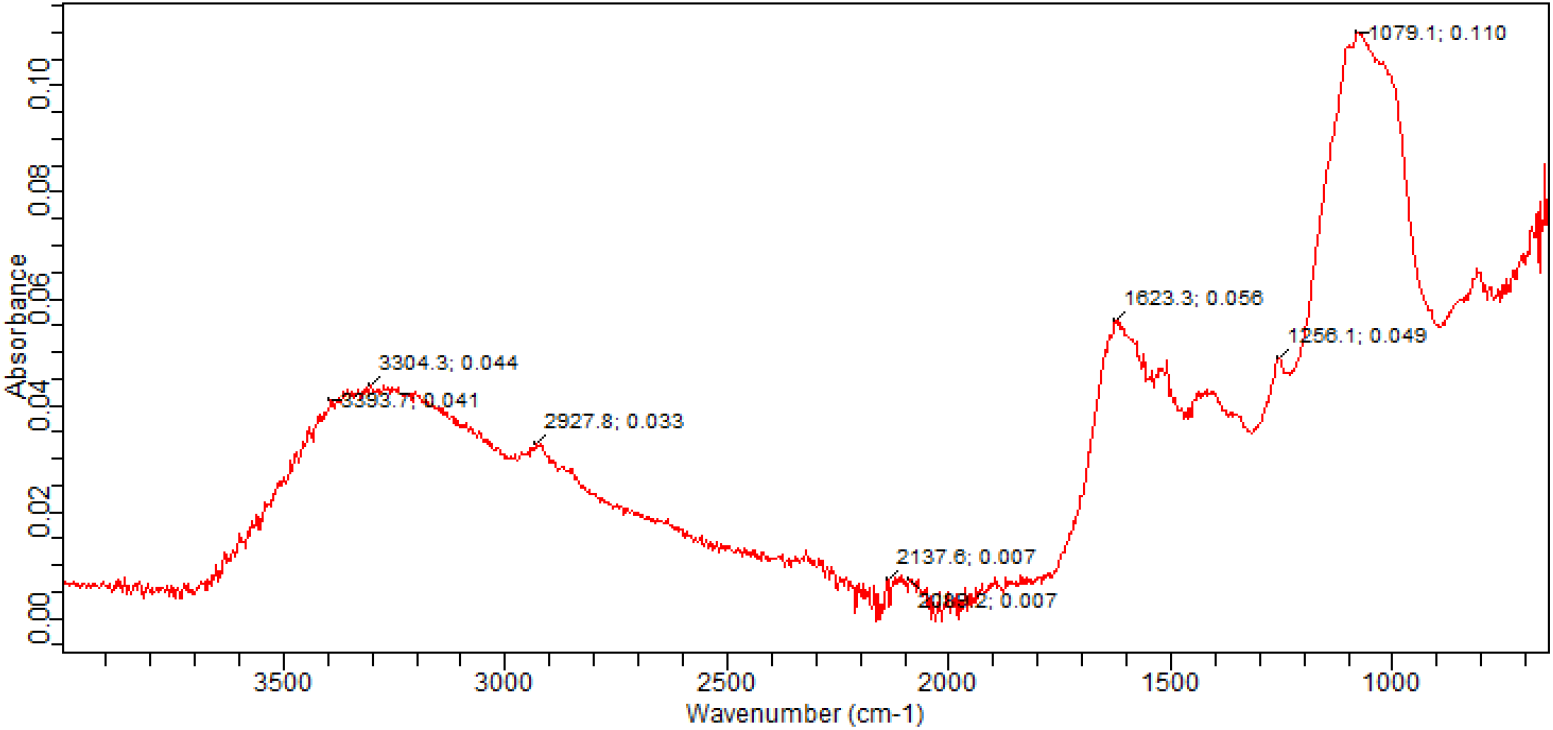
FT-IR spectrum of Mentha Piperita ZnO nanoparticles.

#### Whiteness index assessment of Paint Formulations

The whiteness index of sheets of paints with plant-based additives was measured with a spectrophotometer. The CIE standard Illuminant D65 was used to measure reflective light. Batch 5 to 8 are samples 5 to 8 of our paints and batch 9 is standard biocide paint. This analysis was not performed on paint without biocide as no additive was added to it. [10]

The following factors are measured from the spectrum of whiteness.

❖ DE = total color difference
❖ Da = the difference in the red-green coordinate + Da= marks redder, - Da= marks greener;
❖ Db= the difference in the blue-yellow coordinate + Db= marks yellower, - Db= marks bluer;
❖ DL= the difference in lightness + DL=marks lighter, - DL=marks darker;
❖ DC= the difference in Chroma + DC=marks more brilliant, - DC=marks more opaque.
❖ DH=the difference in color hue
❖ CMC dE= color tolerance

From the data given below, we assess that the plant-based additives affected the whiteness of paint darker in lightness, greener on red-green coordinates, bluer in blue-yellow coordinates, more opaque in chroma, and hue is also affected. dE CMC should be less than 1 for the paint to be acceptable for commercial production. Only sample 6 which is paint with Moringa NPs is close to standard all other samples have affected the whiteness of the paint.

**Table 3:**
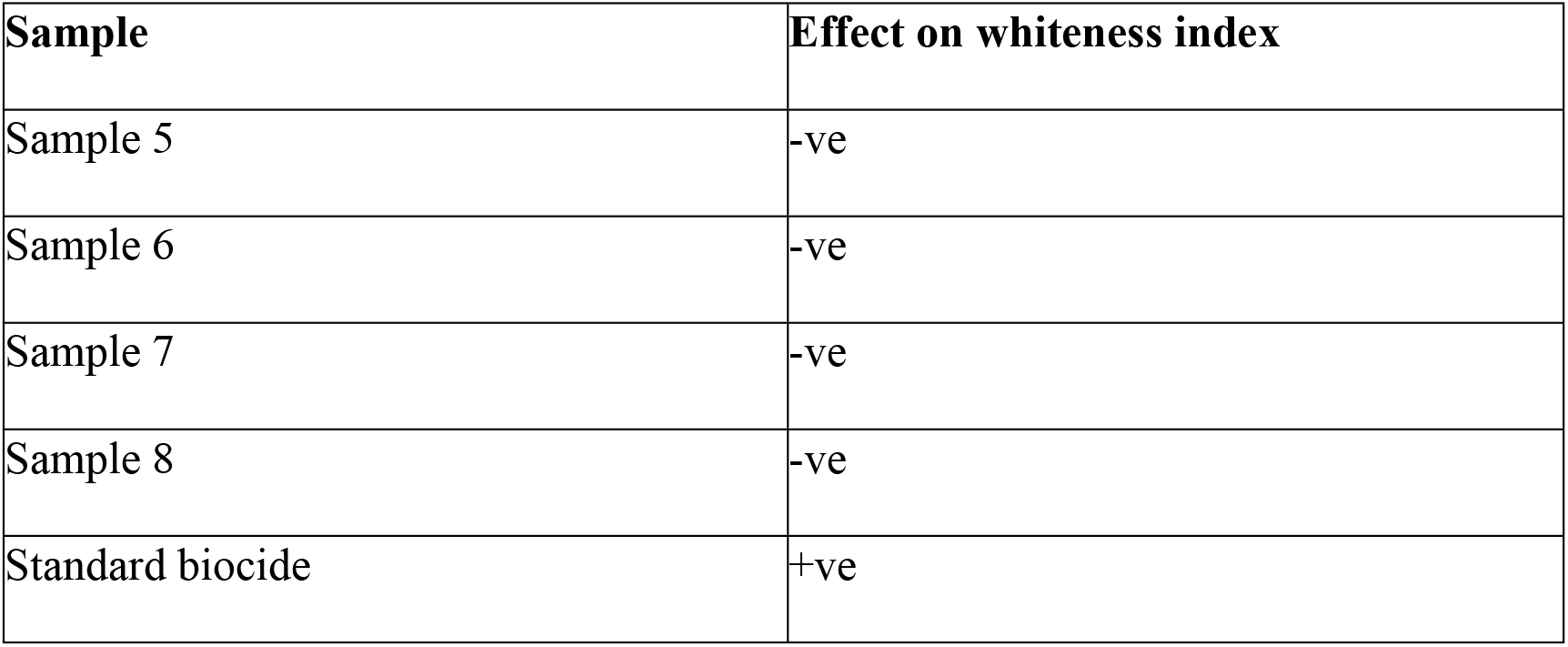
Effect of paint formulation on whiteness index of plain paint.

**Figure 1:**
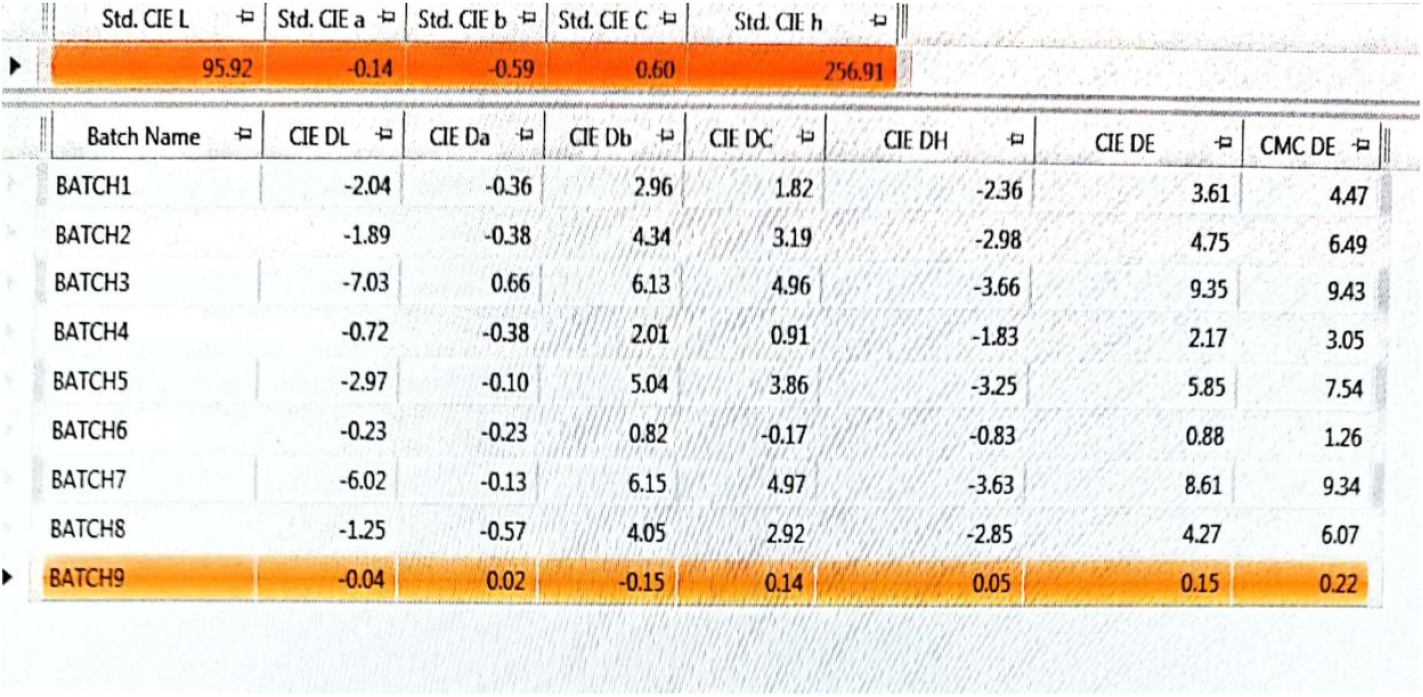
data of whiteness index provided by Diamond Paints industry.

#### 4.5.3 Mosquito repellent activity of Paint formulation

*Aedes aegypti* mature mosquitoes were utilized for the repellency test. Figure 3 shows that paint with a standard biocide is most effective followed by sample 8, sample 6, sample 5, sample 4, and sample without biocide respectively. Thus, we conclude Paint with Peppermint and Moringa extract was most effective than paint with NPs in repelling mosquitos from applied plant material. Biocide showed exceptional results as it is chemically prepared. [2]

**Figure 2:**
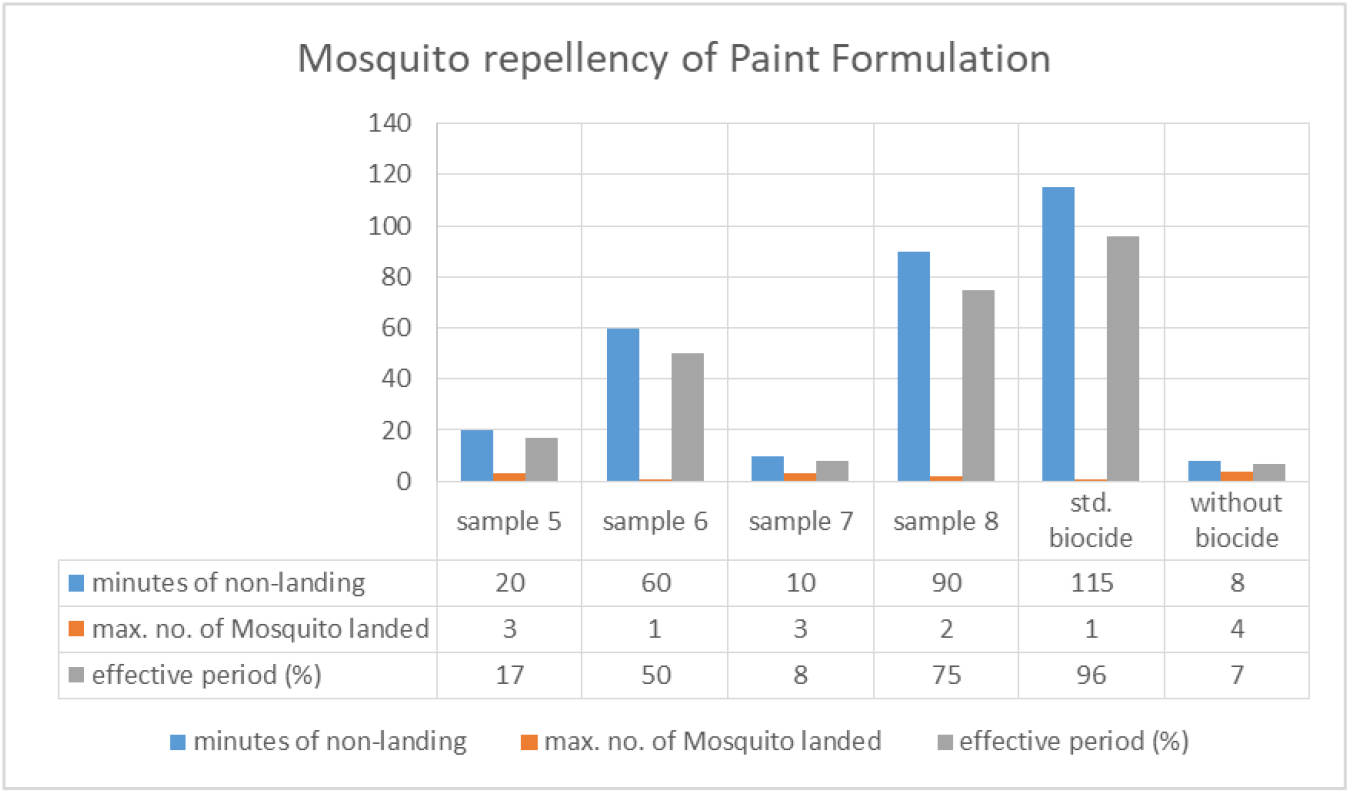
Mosquito repellency of paint formulations.

## Conclusion

It can be concluded from the present research that plants with natural mosquito-repellent properties can be useful in making research coatings that can act as mosquito-repellent as well as antimicrobial. These coatings in addition to being effective also be eco-friendly and less harmful to humans as they are made from plant-based materials. Moringa and Peppermint are easily available plants throughout the year and do not require extensive care. The extract and nanoparticles from these plants are antimicrobial. These can be cultivated at the farm level to produce material for making mosquito-repellent surface coatings. The surface coating prepared using plant material are effective mosquito repellent as proved by the repellency test.

